# Transcriptome Profiling Identifies a Natural Antisense Transcript of IRF4 Associated with EBV Latency

**DOI:** 10.64898/2026.07.24.740669

**Authors:** Ling Wang, Culton R. Hensley, Rifat Jahan, Gifty M. Sarpong, Abigail G. Anderson, Jonathan P. Moorman, Zhi Q. Yao, Shunbin Ning

**Affiliations:** Department of Internal Medicine, Quillen College of Medicine, East Tennessee State University, Johnson City, TN 37614; Center of Excellence for Inflammation, Infectious Diseases and Immunity, Quillen College of Medicine, East Tennessee State University, Johnson City, TN 37614

**Keywords:** lncRNA, IRF4-AS1, EBV, NAT

## Abstract

Epstein-Barr virus (EBV) establishes distinct latency programs that drive B-cell transformation through coordinated viral and host gene regulation. However, the contribution of host long noncoding RNAs (lncRNAs) to EBV-mediated pathogenesis remains poorly understood. Here, we performed transcriptome-wide lncRNA profiling comparing Type I and Type III EBV latency and identified differentially expressed host lncRNAs, among which we characterized a previously unannotated natural antisense transcript (NAT) of IRF4, designated IRF4-AS1. IRF4-AS1 is markedly upregulated in Type III latency and positively correlates with IRF4 expression across EBV-transformed lymphoblastoid cell lines (LCLs) established from diverse clinical backgrounds. Functional studies indicate a context-dependent positive regulatory relationship between IRF4-AS1 and IRF4. While overexpression of IRF4-AS1 increased IRF4 expression and IRF4 overexpression increased IRF4-AS1 levels, loss-of-function analyses suggest that this relationship is modulated by additional regulatory inputs in established transformed cells. In contrast to IRF4, IRF4-AS1 transcription is not regulated by the LMP1- NFκB signaling, indicating that the two transcripts are regulated through partially distinct mechanisms. In addition, we identified MIR3142HG as a highly induced latency-associated lncRNA and found that its embedded miRNAs, miR-146a and miR-3142, exhibited divergent expression patterns, suggesting differential post-transcriptional regulation. Together, these findings identify IRF4-AS1 as a NAT lncRNA for IRF4 and reveal extensive remodeling of host lncRNA networks during EBV latency, providing a foundation for future studies of lncRNA- mediated host-virus interactions in EBV pathogenesis.

**Importance:** EBV drives B-cell transformation through extensive reprogramming of host transcriptional networks, yet the contribution of host long noncoding RNAs (lncRNAs) to this process remains poorly defined. Transcriptome-wide lncRNA profiling, together with functional analysis, identified IRF4-AS1 as a previously unannotated natural antisense transcript of IRF4. IRF4- AS1 is upregulated in Type III latency and positively correlates with IRF4 expression across EBV-transformed cells. Together, this study provides a foundation for future mechanistic studies of lncRNA-mediated host-virus interactions in EBV pathogenesis.

## Introduction

Long noncoding RNAs (lncRNAs) have emerged as an important class of regulatory molecules and potential therapeutic targets in cancer (1–4). The human genome encodes up to 60,000 lncRNA genes, representing roughly 27% of annotated genes (1, 5). These loci include diverse subclasses defined by genomic position, among which natural antisense transcripts (NATs) are widely expressed. Notably, nearly 38% of cancer-associated genomic loci generate sense- antisense transcript pairs, highlighting the extensive involvement of NATs in tumor biology (6). LncRNAs participate in a broad spectrum of cellular processes, and their dysregulation has been linked to many human diseases, including cancer (1, 7–10). However, their roles in viral oncogenesis is just emerging (11, 12).

In contrast to microRNAs, lncRNAs exhibit remarkable functional diversity, acting as molecular scaffolds, modulators of mRNA splicing, stability, or translation, and as decoys for microRNAs or transcription factors (7, 9, 13). In addition, they may function as molecular decoys or “sponges” for miRNAs and transcription factors, thereby modulating gene expression programs (1, 14–18).

Despite increasing recognition of their importance, the contribution of lncRNAs to viral oncogenesis, especially in Epstein-Barr virus (EBV)-associated malignancies, remains incompletely defined. While several EBV-encoded lncRNAs have been studied (2, 19–23), far less is known about how EBV latent infection reprograms host lncRNA networks.

To address this gap, we performed transcriptome-wide lncRNA profiling comparing Type I and Type III EBV latency programs. This analysis uncovered widespread differential expression of host lncRNAs, including a previously unannotated NAT at the IRF4 locus, which we designate IRF4-AS1. IRF4 is a lymphocyte-specific transcription factor with established oncogenic functions in EBV- and Human T-lymphotropic virus type 1 (HTLV1)-infected B/T lymphoma cells (24–35). Given the oncogenic role of IRF4 in these contexts, identifying a latency-associated NAT lncRNA at the IRF4 locus provides an opportunity to uncover new regulatory mechanisms in viral oncogenesis.

## Results

### LncRNA array profiling

To gain insights into the role of lncRNAs in EBV-mediated oncogenesis, we have lncRNA arrays performed by ArrayStar Inc. (a total of 40,173 lncRNA probes and 20,730 mRNA probes) in a pair of isogenic EBV^+^ cell lines, SavI (Type I latency) and its counterpart SavIII (Type III latency). Results identified 2,420 non-redundant lncRNAs (1261 higher and 1159 lower in SavIII vs SavI; cutoff fold: 2.0) (Fig 1, A-B, Suppl. Table 1), including 253 (133 higher and 120 lower) NAT lncRNAs, which are differentially expressed, and most display a correlation pattern with their protein-coding counterpart genes (198 out of 253). Two intergenic lncRNAs, MIR3142HG and MIR155HG (as known as B cell integration cluster (BIC)), are 1422-fold and 9-fold higher, respectively, in SavIII vs SavI cells (Fig 1, C). MIR3142HG harbors miR-146a and miR-3142, and MIR155HG harbors miR-155. miR-146a and miR-155 are well documented to be deregulated by EBV latent infection (36).

**Fig 1.**
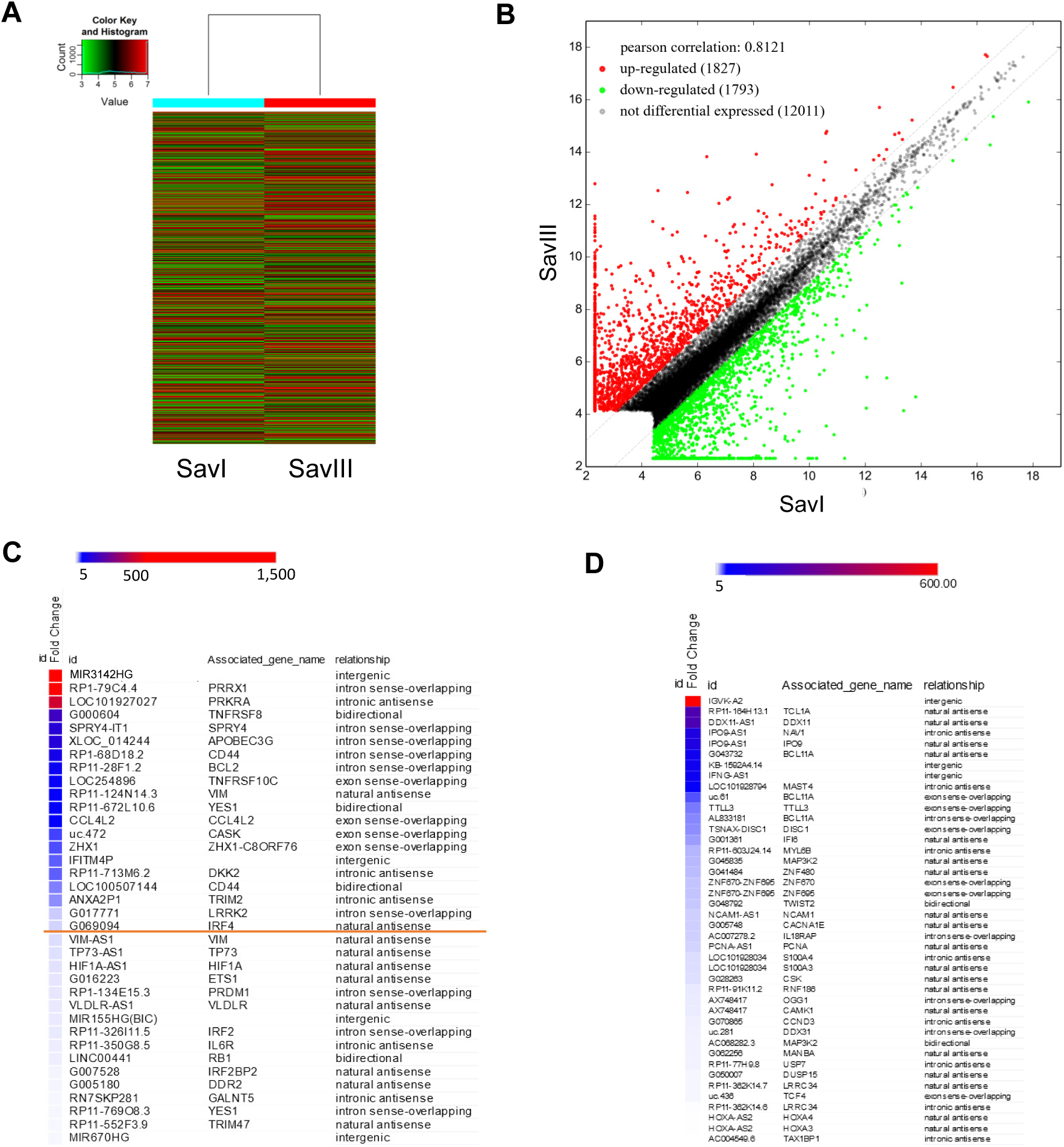
lncRNA array profiling. **A.** Hierarchical clustering of differentially expressed lncRNAs. **B**. Scatter plot of differentially expressed lncRNAs. **C-D**. Selected upregulated (C) and downregulated (D) lncRNAs in SavIII vs SavI (cutoff: >5 fold) relevant to our research interests are shown, including miR-146 and miR-155 host genes (miR3142HG and miR155HG/BIC).

In this study, we focus on the category of NAT lncRNAs whose differential expression was detected in our high throughput assays (Fig 1, C-D). Interesting NAT lncRNAs that are significant higher in SavIII vs SavI cells with potential roles in EBV oncogenesis include G069094 (NAT for IRF4, 13-fold), VIM-AS1 (11-fold), TP73-AS1 (11-fold), HIF1A-AS1 (10-fold), G007528 (NAT for IRF2BP2, 7-fold), and G005180 (NAT for DDR2, 6.8-fold) (Fig 1, C).

### IRF4-AS1 as an NAT lncRNA for IRF4

We found that the transcript G069094 derived from chromosome 6 overlaps with IRF4 mRNA in sequence. Specifically, the mature G069094 transcript has 2,518 nt in length without a poly(A) tail (not shown), and the region spanning 1116-1184 overlaps with 5’-end UTR (head-to-head) of IRF4 mRNA (NM_002460, 1-68) (Fig 2, A-B). Subcellular fraction qPCR results show that IRF4-AS1, like IRF4, is localized in both the nucleus and cytoplasm (Fig 2, C). Based on these findings, we conclude that the transcript G069094 is a NAT lncRNA of IRF4. We thus re-named it IRF4-AS1.

**Fig 2.**
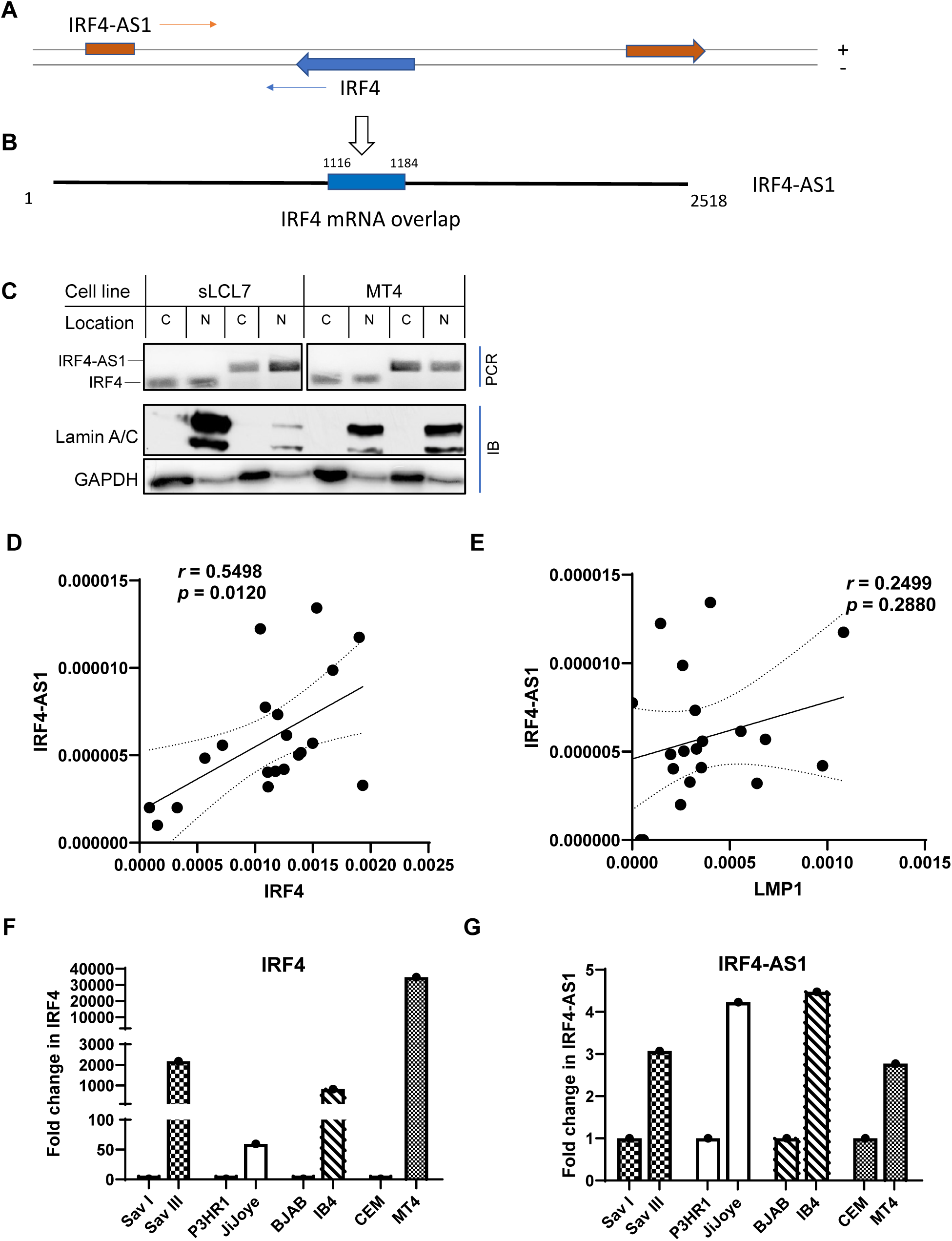
Identification of IRF4-AS1 A-B. A scheme of IRF4/IRF4-AS1 gene locus (A) and IRF4-AS1 mature transcript (B). The mature IRF4-AS1 has 2,518 bp in length, and the region spanning 1116-1184 overlaps with 5’- end UTR of IRF4 mRNA. **C**. RT-PCR analysis shows that IRF4-AS1 and IRF4 are localized in both the cytoplasm and nucleus. The cytoplasmic marker GAPDH and the nuclear marker Lamin A/C were probed by immunoblotting. C: Cytoplasmic. N: Nuclear. **D-E.** Pearson correlation analysis for IRF4-AS1 with IRF4 and with LMP1 mRNA levels across LCLs, as determined by qPCR. **F-G**. Evaluation of IRF4 and IRF4-AS1 transcript levels in viral latency programs by qPCR. Representative results from at least three independent repeats are shown. Error bars represent mean ± SE of triplicates. IRF4 mRNA and IRF4-AS1 transcript levels in SavI, P3HR1, BJAB, and CEM cells were set to 1, and those in the corresponding counterpart cell lines are shown as fold changes.

We validated IRF4:IRF4-AS1 expression pattern using quantitative real-time PCR (qPCR) across a panel of in vitro transformed lymphoblastoid cell lines (LCLs) from healthy subjects and patients with Human immunodeficiency virus-1 (HIV1) infection, and spontaneous LCLs derived from patients with infectious mononucleosis (IM) and from transplant recipients (37, 38). Pearson correlation analysis shows that the expression levels of IRF4 and IRF4-AS1 were positively correlated across these EBV-transformed LCLs (r = 0.55, p = 0.012). (Fig 2, D). In contrast, LMP1 and IRF4-AS1 are not significantly correlated (r=0.2499, p=0.2880) (Fig 2, E).

We further evaluated both IRF4 and IRF4-AS1 transcript expression in viral latency programs by qPCR. The results show that both IRF4 mRNA and IRF4-AS1 transcript levels are higher in EBV Type III latency compared to Type I latency, and high in HTLV1-infected cells compared to HTLV1-negative cells (Fig 2, F-G). These findings suggest that IRF4-AS1 and its sense partner, IRF4, may share a common transcriptional regulatory mechanism.

### MIR3142HG

Another relatively novel lncRNA differentially expressed between SavIII and SavI in our array is MIR3142HG, which harbors two miRNA genes encoding miR-3142 and miR-146a (Fig 3, A). qPCR analysis showed that MIR3142HG expression correlates with EBV latency programs (Fig 3, B). Furthermore, MIR3142HG expression positively correlated with both IRF4 (r = 0.5293, p = 0.0164) and LMP1 (r = 0.5699, p = 0.0087) across EBV-transformed LCLs (Fig 3, C–D). These findings suggest that MIR3142HG is associated with activation of the LMP1-IRF4 signaling axis during EBV latency.

**Fig 3.**
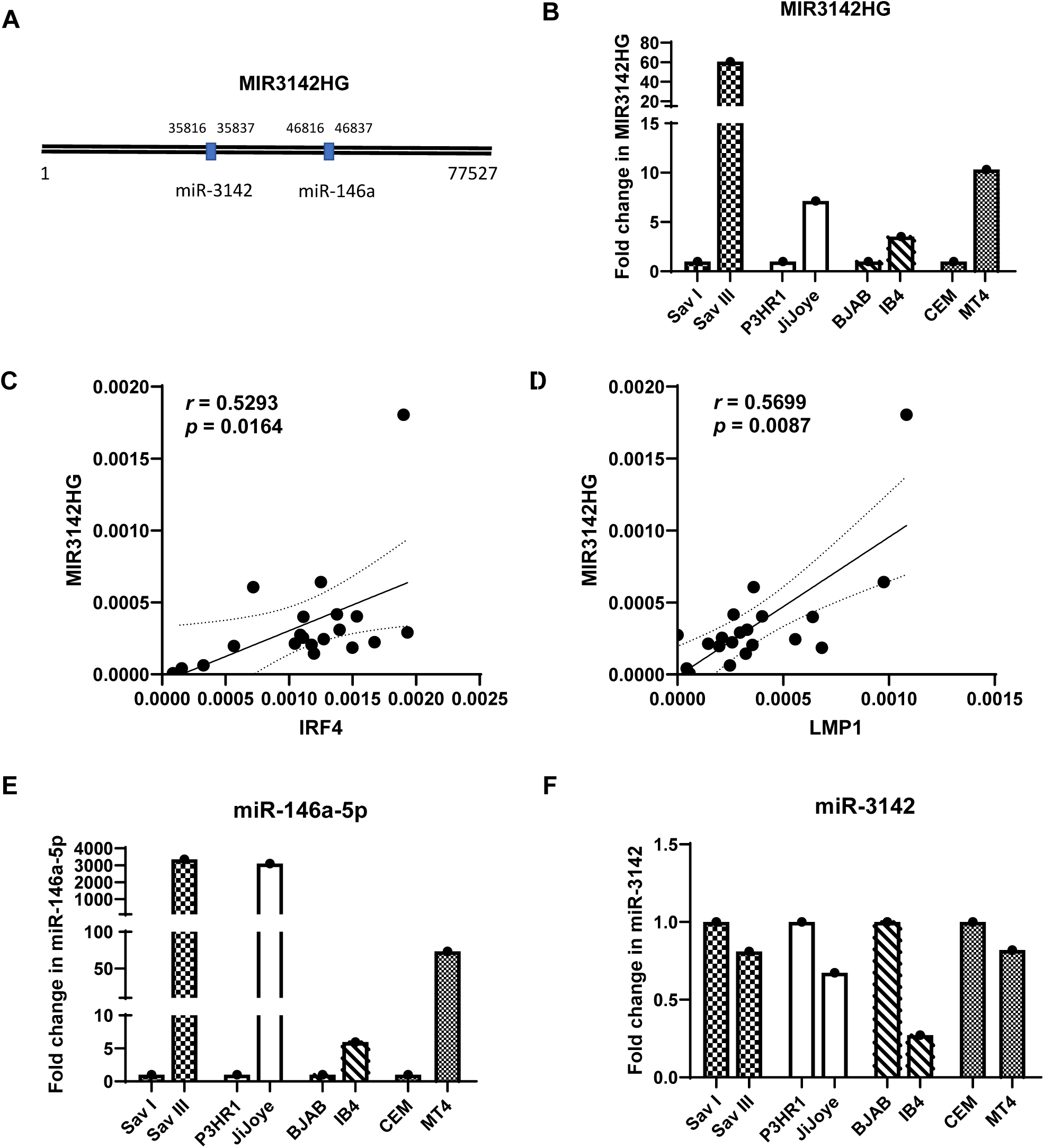
Characterization of MIR3142HG. **A**. Schematic diagram of the MIR3142HG gene locus. **B**. Correlation of MIR3142HG transcript levels with viral latency programs. **C-D**. Spearman correlation analysis of MIR3142HG transcript levels with IRF4 (C) and LMP1 mRNA levels (D) across LCLs, as determined by qPCR. **E-F**. Correlation analysis for the MIR3142HG-derived miRNAs, miR-146a-5p (E) and miR-3142 expression levels with viral latency programs (F), as determined by qPCR. Representative results from at least three independent experiments are shown as mean ± SE of triplicates. In B, E, and F, expression levels in SavI, P3HR1, BJAB, and CEM cells were set to 1, and those in the corresponding counterpart cell lines are shown as fold changes.

miR-146a-5p exhibited an expression pattern similar to its host transcript MIR3142HG (Fig 3, E). In contrast, miR-3142 showed an inverse expression pattern (Fig 3, F). This divergence suggests that miR-146a-5p and miR-3142, despite being encoded within the same primary transcript, may be differentially regulated at the post-transcriptional level, thereby decoupling mature miRNA abundance from host transcript expression during EBV latency.

### IRF4-AS1 and IRF4 reciprocal regulation

We chose IRF4-AS1 for further study because we have shown that IRF4 is induced and activated by LMP1 (28, 29), and is critical for survival and proliferation of EBV- or HTLV1- positive lymphoma cells and other blood cancer cells (34, 39, 40). We also included MIR3142HG for comparative analysis, as one of its encoded products, miR-146a, has been extensively studied in the context of EBV latency and immune regulation (41, 42).

In general, NAT lncRNAs and their overlapping sense protein-coding genes often form coupled regulatory loci. The sense gene locus can influence antisense lncRNA expression through shared regulatory elements, transcriptional coupling, chromatin effects, or RNA- mediated mechanisms. Conversely, NAT lncRNAs can regulate the expression of their overlapping sense genes in *cis* (so-called *cis*-NATs), via transcriptional interference, chromatin remodeling, and RNA-RNA interactions, highlighting bidirectional regulation at the same genomic locus (43–48).

To test if IRF4 regulates IRF4-AS1 expression, we transiently expressed Flag-IRF4 in EBV- negative BJAB cells that do not express detectable IRF4, and quantitated IRF4-AS1 expression by qPCR after 2 days post nucleofection. Results show that IRF4 overexpression (OE) significantly increases IRF4-AS1 and MIR3142HG transcript levels, but not MALAT1 levels that can serve as a negative control in this experiment (Fig 4, A). These results are consistent with their correlations across EBV-transformed LCLs (Fig 2, D; Fig 3, C).

**Fig 4.**
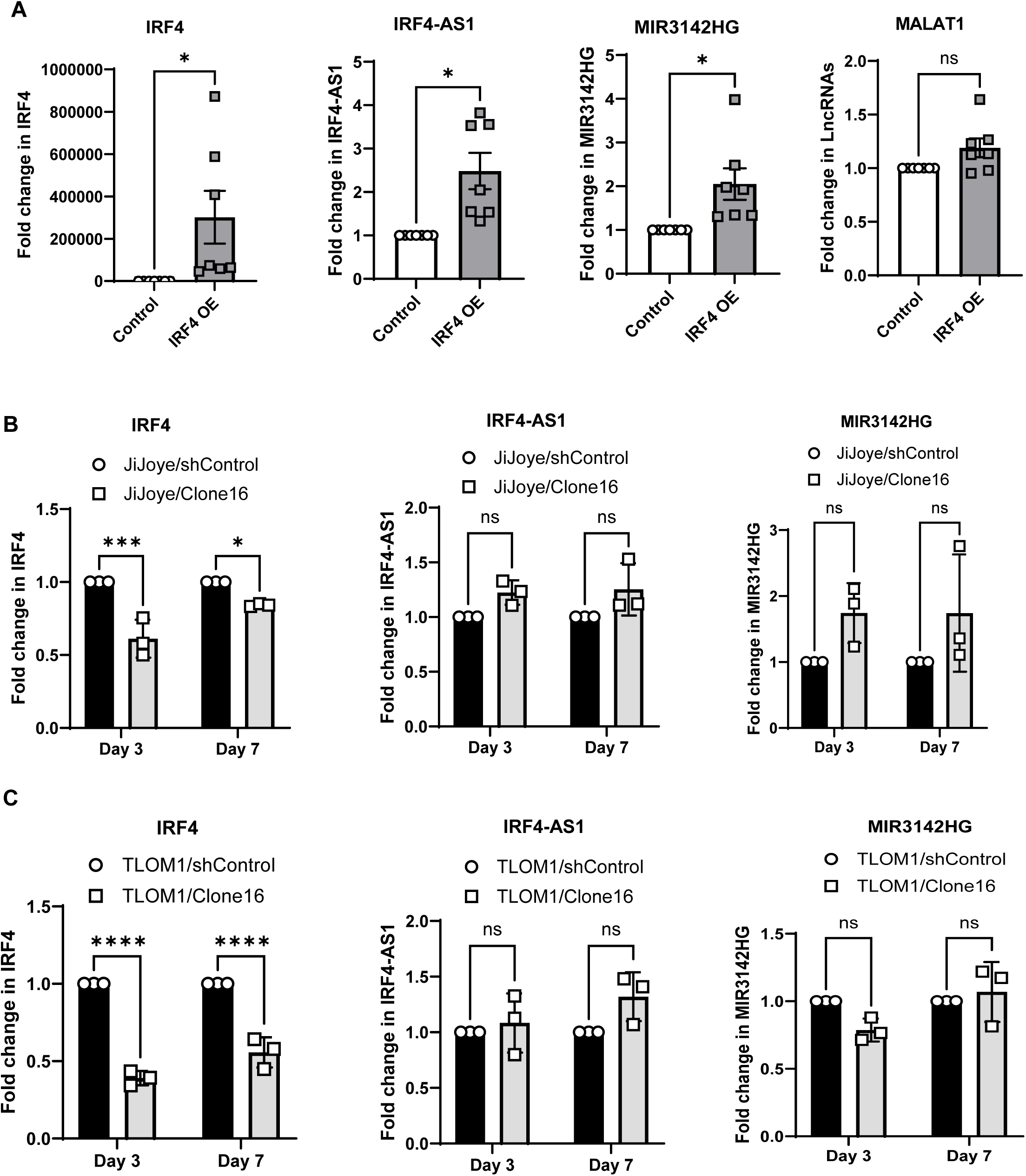
Regulation of IRF4-AS1 transcription by IRF4. **A**. Transient expression of IRF4 in BJAB cells upregulates the transcript levels of IRF4-AS1 and MIR3142HG, but not those of MALAT1. Summary data from seven independent experiments are presented, with error bars representing the mean ± SE. **B-C**. shRNA-mediated knockdown of IRF4 in JiJoye and MT4 cells did not downregulate transcript levels of IRF4-AS1 and MIR3142HG. Summary data from three independent experiments are presented, with error bars representing the mean ± SE. The IRF4 mRNA levels, IRF4-AS1, MIR3142HG and MALAT1 transcript levels in BJAB cells expressing control, as well as JiJoye and IB4 cells expressing Control shRNA (shControl) were set to 1, and those in the corresponding counterpart cell lines are shown as fold changes. Statistical significance was determined by a paired t-test (A) or two-way ANOVA (B-C), and is shown as p<0.05 (*), p<0.01 (**), <0.001 (***), or <0.0001 (****). n.s.: not significant.

We then evaluated IRF4-AS1 expression in JiJoye and TLOM1 cells stably expressing IRF4 shRNA, which were established in our previous study (30). Surprisingly, we did not detect significant downregulation of IRF4-AS1 in either IRF4-deficient cell lines (Fig 4, B-C).

To determine if IRF4-AS1 regulates IRF4 expression, we constructed a pcDNA3.1-IRF4- AS1 expression plasmid, and transfected it into BJAB cells. Results show that IRF4-AS1 overexpression significantly increases endogenous IRF4 mRNA level. Interestingly, IRF4-AS1 overexpression also increases transcription of MIR3142HG (Fig 5, A).

**Fig 5.**
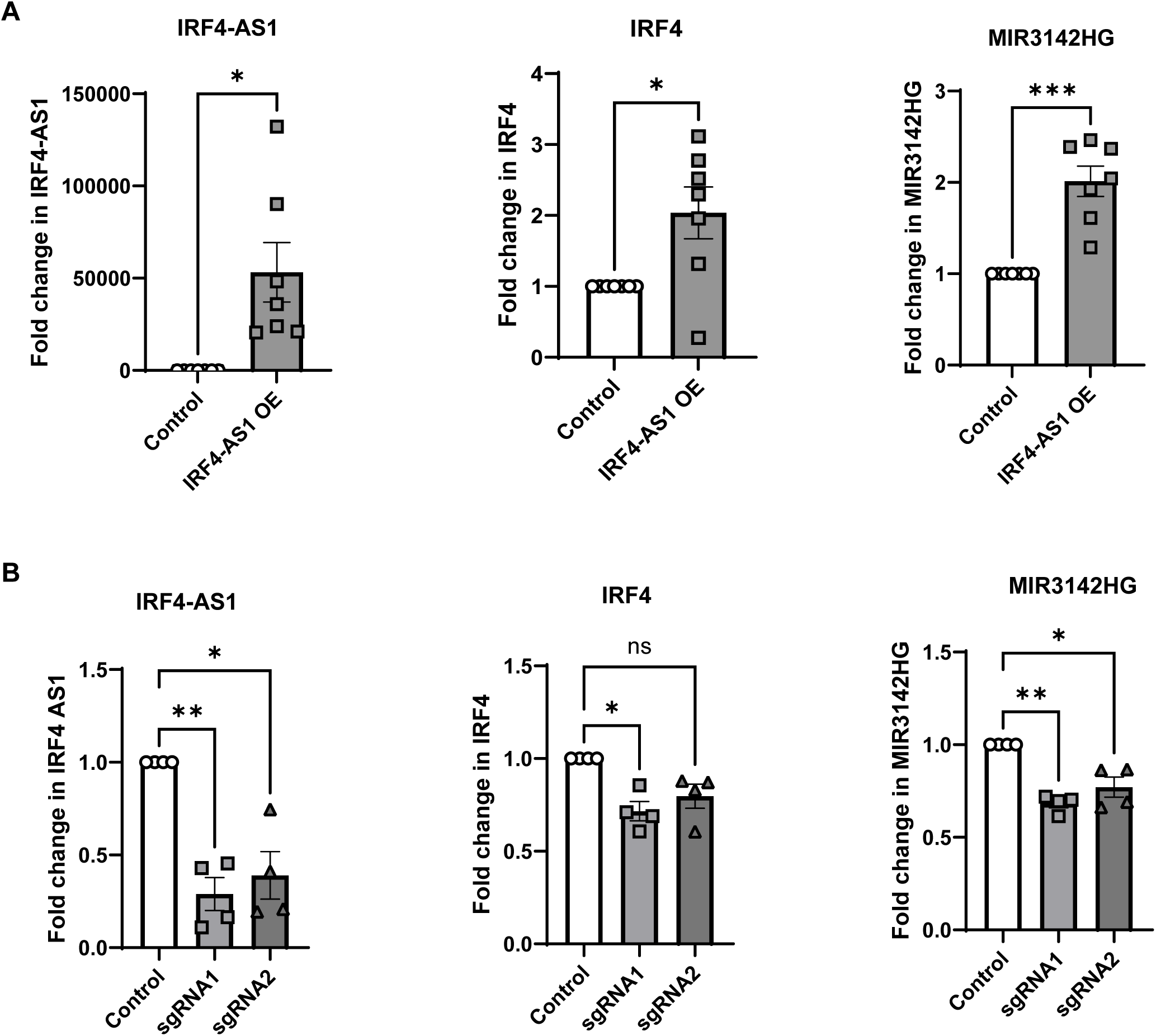
Regulation of IRF4 transcription by IRF4-AS1. **A**. Transient expression of IRF4-AS1 in BJAB cells upregulates the transcript levels of IRF4 and MIR3142HG. Summary data from seven independent experiments are presented, with error bars representing the mean ± SE. **B**. sgRNA-mediated knockdown of IRF4-AS1 in JiJoye cells downregulates the transcript levels of IRF4 and MIR3142HG. Summary data from four independent experiments are presented, with error bars representing the mean ± SE. IRF4 mRNA levels, IRF4-AS1 and MIR3142HG transcript levels in BJAB cells expressing control, as well as JiJoye cells expressing Control sgRNA were set to 1, and those in the corresponding counterpart cell lines are shown as fold changes. Statistical significance was determined by a paired t-test (A) or one-way ANOVA (B), and is shown as p<0.05 (*), p<0.01 (**), <0.001 (***), or <0.0001 (****). n.s.: not significant.

Next, we have constructed two IRF4-AS1 sgRNAs (or control sgRNA) into pLenti- CRISPRv2, and used them to generate JiJoye stable cell lines. qPCR results show that IRF4- AS1 depletion modestly reduced IRF4 expression by day 3, with sgRNA1 producing a statistically significant effect (∼30%, p<0.05), whereas MIR3142HG expression was consistently reduced by both IRF4-AS1 sgRNAs (Fig 5, B).

Together, the transient overexpression experiments suggest that IRF4 and IRF4-AS1 participate in a reciprocal positive regulatory relationship. However, this relationship was not fully recapitulated in loss-of-function experiments in virus-transformed cell lines. Specifically, IRF4 knockdown did not significantly reduce IRF4-AS1 expression, whereas IRF4-AS1 depletion produced only a modest reduction in IRF4 expression, suggesting that additional regulatory mechanisms maintain expression of both transcripts in established transformed cells. Consistent with this interpretation, we and others have shown that IRF4 is strongly induced by the viral oncoproteins LMP1 and Tax in EBV- and HTLV1-transformed cells, respectively (24–26, 30). These viral regulatory pathways may compensate for or mask the regulatory interactions observed under transient expression conditions.

### Regulation of IRF4-AS1 and MIR3142HG by EBV LMP1 signaling

*Cis*-NATs are typically RNA polymerase II-dependent RNA transcripts that utilize the same machinery as their sense partners for transcription (17, 44, 49). In this regard, we tested if IRF4-AS1 transcription is regulated by LMP1-NF-κB axis, since IRF4 is induced by this pathway (24).

First, we transfected LMP1 expression plasmids into BJAB cells. qPCR results show LMP1 significantly upregulates both IRF4-AS1 and MIR3142HG transcripts (Fig 6, A).

**Fig 6.**
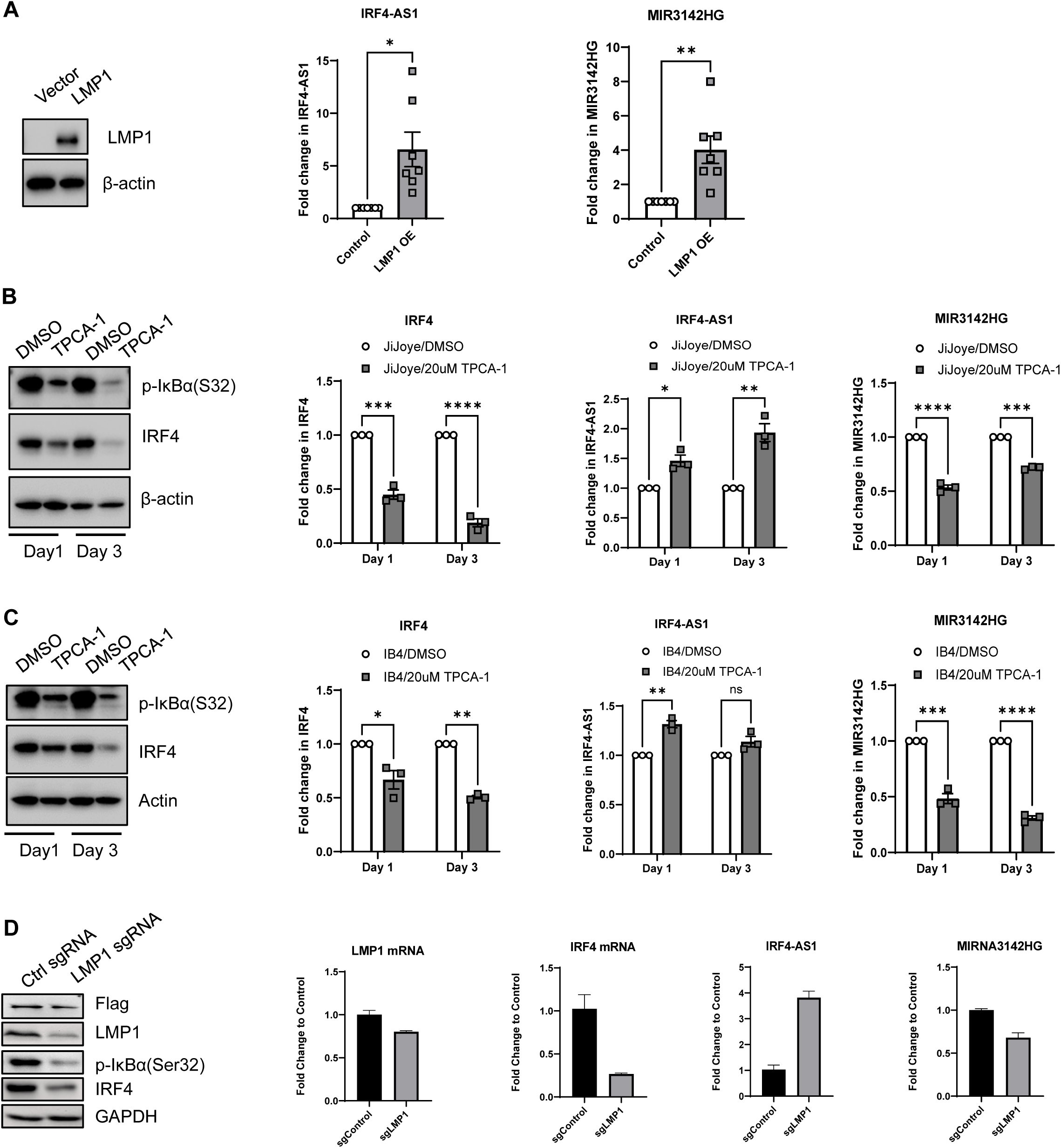
IRF4 regulates IRF4-AS1 transcription. **A.** LMP1 transient expression in BJAB cells upregulates the transcript levels of IRF4-AS1 and MIR3142HG. Summary data from seven independent experiments are presented, with error bars representing the mean ± SE. **B-C**. TPCA-1 treatment in JiJoye (B) and IB4 (C) cells downregulates the transcript levels of MIR3142HG, but not IRF4-AS1. Summary data from three independent experiments are presented, with error bars representing the mean ± SE. Statistical significance was determined by a paired t-test (A) or two-way ANOVA (B-C), and is shown as p<0.05 (*), p<0.01 (**), <0.001 (***), or <0.0001 (****). n.s.: not significant. **D**. sgRNA- mediated LMP1 depletion in IB4 cells downregulates MIR3142HG, but upregulates IRF4-AS1 transcript levels. Representative results with mean ± SE of triplicates from three independent experiments are shown. IRF4 mRNA levels, IRF4-AS1 and MIR3142HG transcript levels in BJAB cells expressing control, JiJoye and IB4 cells treated with DMSO, and JiJoye cells expressing Control sgRNA were all set to 1, and those in the corresponding counterpart cell lines are shown as fold changes.

We then inhibited NFκB activity by the IKKβ-specific inhibitor TPCA-1 in JiJoye and IB4 cells. TPCA-1 treatment remarkably inhibits phosphorylation of IκBα(S32), an indicator of NFκB activity. Results show that IRF4 and MIR3142HG mRNA levels were significantly downregulated by TPCA-1 treatment. Surprisingly, IRF4-AS1 mRNA levels were slightly upregulated (Fig 6, B-C). Consistently, LMP1 depletion in IB4 cells downregulates MIR3142HG (and IRF4) transcript, but upregulates IRF4-AS1 transcript (Fig 6, D).

Together with Fig 2E and Fig 3D, these results suggest that MIR3142HG, but not IRF4-AS1, is directly regulated by the LMP1-NFκB axis.

## Discussion

EBV latent infection extensively reprograms host transcriptional networks to sustain proliferation and survival. Although EBV-encoded viral lncRNAs, e.g. BARTs and BHLF1 lncRNA, have been investigated for years (19–22), the contribution of host lncRNAs to EBV- driven pathogenesis remains poorly defined (11). In this study, transcriptome-wide profiling from a pair of EBV-infected isogenic cell lines revealed widespread dysregulation of host lncRNAs between latency I and III programs, including a substantial subset of NAT lncRNAs. Notably, the majority of differentially expressed NAT lncRNAs exhibited coordinated expression with their corresponding protein-coding genes, suggesting that antisense transcription represents a common feature of host transcriptional reprogramming during EBV latency.

A principal finding of this study is the identification of IRF4-AS1, a previously unannotated NAT lncRNA located at the IRF4 locus. NAT lncRNAs have emerged as important regulators of gene expression through mechanisms such as transcriptional interference, chromatin remodeling, and RNA-mediated regulation (43–49). Consistent with these paradigms, IRF4- AS1 expression positively correlates with that of its sense partner IRF4 across multiple virus- transformed cell systems. However, functional analyses indicate that the IRF4-AS1/IRF4 relationship is more complex than a simple positive-feedback model.

Although transient expression of LMP1 induced both IRF4 and IRF4-AS1 expression, inhibition of canonical NFκB activity or LMP1 depletion in EBV latency III reduced IRF4 expression but upregulated IRF4-AS1. These observations suggest that coordinated expression of IRF4 and IRF4-AS1 does not require identical upstream regulatory mechanisms. Instead, multiple transcriptional pathways may converge on the IRF4 locus, resulting in coordinated but not necessarily equivalent regulation of the sense and antisense transcripts.

In addition to IRF4-AS1, this work identified MIR3142HG as one of the most highly induced lncRNAs associated with latency III. Interestingly, MIR3142HG and LMP1 transcripts correlate across EBV LCLs and EBV latency programs, and gain- and loss-of-function analyses consistently indicate that MIR3142HG is induced by the LMP1-NFκB signaling axis, and that IRF4-AS1 may contributes to its transcription.

Collectively, these observations support a model in which EBV latency establishes a multilayered regulatory network integrating transcriptional and post-transcriptional mechanisms, in which sense-antisense gene pairs and miRNA-host gene transcripts are coordinately induced yet differentially regulated through distinct signaling and buffering pathways.

In contrast to the MIR3142HG product miR-146a, which has been extensively investigated in immunity, inflammation, and cancer (50), little is known about the biological functions and regulatory mechanisms of MIR3142HG itself and the other product miR-3142. While one study identified that MIR3142HG is strongly linked to NFκB-driven inflammatory programs implicated in pathogenesis of systemic lupus erythematosus (SLE) (51), another study shows that the product miR-3142 functions as an oncogenic miRNA that promotes Akt activation and chemoresistance in chronic myeloid leukemia (52). Our results suggest that MIR3142HG may also play a role in EBV-driven pathogenesis by acting as a downstream target of the LMP1- NFκB signaling.

The molecular basis of IRF4-AS1 and IRF4 co-regulation remains to be defined, and genome-wide targets of IRF4-AS1 have not yet been characterized. Future transcriptomic and mechanistic studies will be required to determine whether IRF4-AS1, MIR3142HG, and miR- 3142 functionally contribute to EBV-associated pathogenesis.

From a therapeutic perspective, lncRNAs embedded within EBV-driven transcriptional networks may represent potential targets for RNA-based intervention. However, the context- dependent nature of NAT-sense gene relationships highlighted in this study cautions that perturbation of a NAT lncRNA may not produce a predictable or proportional effect on its cognate protein-coding gene, particularly in virus-transformed cellular contexts where compensatory regulatory mechanisms are likely present. Rigorous functional validation will therefore be essential to determine whether these transcripts can be reliably exploited for therapeutic modulation.

In summary, this work identifies distinct regulatory behaviors of host lncRNAs during EBV latency. IRF4-AS1 is regulated through mechanisms that are partially uncoupled from canonical NFκB signaling despite correlation with IRF4, whereas distinct approaches consistently demonstrated MIR3142HG as a downstream transcriptional target of the LMP1-NFκB signaling axis. In addition, MIR3142HG-derived miR-146a and miR-3142 exhibit divergent expression patterns, indicating differential processing during microRNA maturation from host lncRNA expression. Together, these findings support a multilayered regulatory architecture in which EBV latency reshapes host lncRNA networks through both transcriptional control and post- transcriptional processing, providing a framework for future studies of lncRNA-mediated host- virus interactions in EBV latency and pathogenesis.

## Materials and Methods

### Cell Lines

The paired cell lines, SavI and SavIII, and P3HR1 and JiJoye were derived from EBV- positive human Burkitt’s lymphoma (BL) patients. P3HR1 lacks LMP1 expression due to the absence of the entire EBNA2 ORF (53). JiJoye-shCtrl and JiJoye-shIRF4 stable cell lines were used in our previous publication (30). BJAB is an EBV-negative BL line. The lymphoblastic cell line (LCL) IB4 was established from umbilical cord B-lymphocytes latently transformed with EBV *in vitro*. CEM (HTLV1-) and MT4 (HTLV1+) are T cell lines derived from leukemia patients. Spontaneous LCLs derived from EBV-positive patients, including infectious mononucleosis (IM; 4 patients), transplant recipients (4 patients), or post-transplant lymphoproliferative disorder (1 patient) were generous gifts from Dr. Delecluse, and details are provided in relevant publications (37, 38).

B and T cell lines are cultured with RPMI1640 medium plus 10% heat inactivated Fetal Bovine Serum (FBS, Gibco, Jenks, OK) and antibiotics (Life Technologies, Carlsbad, CA).

### Human subjects

The study protocol for collecting whole blood from people living with HIV (PLHIV) was approved by the joint Institutional Review Board (IRB) of East Tennessee State University and the James H. Quillen VA Medical Center (ETSU/VA IRB #0519.24s). Written informed consent was obtained from all participants. Whole blood from healthy subjects was obtained from BioIVT Elevating Science (Gray, TN). Subjects with malignancy, transplantation, HBV or HCV infection, or receiving immunosuppressive drug treatment were excluded. Peripheral blood mononuclear cells (PBMCs) were isolated from fresh heparinized whole blood using SepMate- 50 (STEMCELL Technologies, Vancouver, Canada) and cryopreserved in liquid nitrogen until use. The characteristics of the human subjects are shown in Table 1.

**Table 1.**
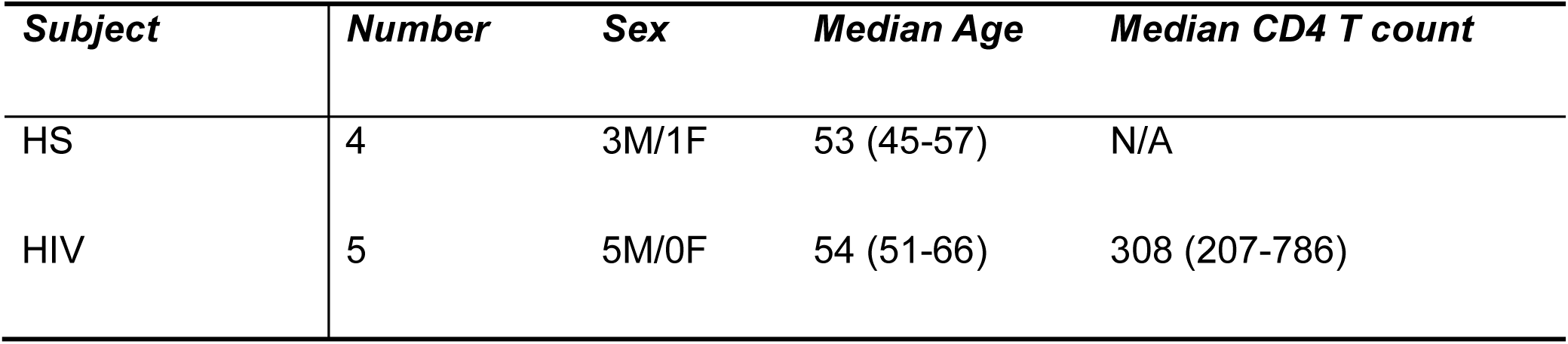
The characteristics of the human subjects.

### Establishment of EBV-transformed lymphoblastoid cell lines (LCLs)

The EBV-positive B95.8 monkey B cell line was stimulated with 500 ng/mL of 12-O- tetradecanoylphorbol 13-acetate (TPA, Sigma, Burlington, MA, USA) for 3 h, followed by washing three times with Dulbecco’s phosphate-buffered saline (DPBS, Gibco). The cells were subsequently resuspended in fresh complete RPMI1640 medium and cultured for 5 days, at which point the supernatant was harvested. PBMCs were obtained from 5 PLWH on ART with undetectable viremia (HIV-RNA <20 copies/mL) and 4 healthy subjects, and seeded in 12-well plates (3∼4 × 10^6^ cells each well), and then exposed to approximately 25 µL of EBV virions per well in the presence of 200 ng/mL of cyclosporin A (CSA, Sigma, Saint Louis, MO, USA). Once visible clusters appear, expand cultures into larger vessels, and established LCLs can be maintained long-term with regular passaging.

In a separate experiment, PBMCs were depleted of CD4⁺ T cells using human CD4 MicroBeads (Miltenyi Biotec, Auburn, CA, USA), then seeded into 24-well plates at ∼1.5 × 10^6^ cells per well and infected with 20 µl of EBV per well. Cells were cultured in RPMI 1640 medium and harvested 35 days post-infection.

### lncRNA array profiling

Total RNA was extracted from SavI and SavIII cells using the RNeasy mini kit (Qiagen). lncRNA array and bioinformatics data processing were performed by ArrayStar Inc., using the Human V5.0 LncRNA Array Service with the chip containing a total of 39,317 lncRNA and 21,174 mRNA probes.

### Cellular fractionation and lncRNA extraction

Nuclear and cytoplasmic fractionation was performed with the NE-PER™ Nuclear and Cytoplasmic Extraction Reagents (TheromoFisher Scientific), following the manufacturer’s instructions. Click-iT® Nascent RNA Capture Kit (Invitrogen) was used to extract nascent RNAs from nuclear and cytoplasmic fractions, following the manufacturer’s instructions.

### DNA constructs

IRF4-AS1 cDNA (2.5 kb) was synthesized and cloned into pcDNA3.1(+) by GeneArt (San Diego, CA). IRF4-AS1 sgRNAs (#1: 5’-GCGATGTTCTCTAAACACCG-3’; #2: CGTTCACAAACATACCAGAG) were cloned into pLenti-CRISPRv2 eSpCas9 by GenScript (Piscataway, NJ). Flag-IRF4 and LMP1 expression plasmids were used in our previous publications (28, 30). LMP1 and control sgRNA cloned in pLenti-Guide-Puro were gifts from Dr. Gewurz (54).

### Transfection

Cell lines were transfected using the GenePulser XCell (Bio-Rad, Hercules, CA), with optimized programs, or nucleofected with a 4D-nucleofector using the P3 Primary Cell 4D- Nucleofector® X Kit L (Lonza, Basel, Switzerland, transduced with lentivirus as detailed in our publications (30–32).

### TPCA-1 drug treatment

TPCA-1, an IKKβ-specific inhibitor, was purchased from MedChemExpress LLC (Monmouth Junction, NJ). JiJoye and IB4 Cell were treated with 20 µM TPCA-1, and cells were collected at 1 day and 3 days post-treatment for Western blotting and qPCR analyses.

### CRISPR/Cas9-mediated depletion

Plasmids harboring IRF4-AS1 or control sgRNAs in pLenti-CRISPRv2 eSpCas9 were introduced into cells by lentivirus-mediated transfection. Lentivirus packing, production, and infection were detailed in our previous publication (30, 55, 56). Stable transfectants were selected with 1 µg/ml puromycin for 2 weeks. Expression of sgRNA is induced by 1 µg/ml doxycycline (Sigma). For CRISPR/Cas9-mediated depletion of LMP1, a protocol from Dr. Gewurz’s lab was adopted (57). Briefly, IB4 cells stably expressing Cas9 were established with 10 mg/ml Blasticidin selection, and then infected with lentiviruses harboring LMP1 or control sgRNA. Cells were selected with 1.0 µg/ml puromycin for 2 days and collected before massive cell death.

### Western blotting

Cells were lysed on ice in RIPA buffer (Boston BioProducts, Ashland, MA) supplemented with a protease inhibitor cocktail (Roche Diagnostics GmbH, Mannheim, Germany). Protein concentrations were determined using the Coomassie (Bradford) Protein Assay (Bio-Rad, Hercules, CA). Equal amounts of protein were separated by SDS-PAGE and transferred onto nitrocellulose membranes (MilliporeSigma, Burlington, MA). Membranes were blocked with 5% nonfat milk in Tris-buffered saline containing 0.1% Tween-20 (TBST) and incubated with primary antibodies against Lamin A/C, GAPDH (Santa Cruz Technologies, Santa Cruz, CA), phospho-IκBα (Ser32), anti-β-Actin (Cell Signaling Technology; Danvers, MA), IRF4 (ProteinTech, Rosemont, IL) and LMP1 (MilliporeSigma). After incubation with the appropriate horseradish peroxidase (HRP)-conjugated secondary antibodies (Cell Signaling Technology), immunoreactive proteins were detected using Amersham ECL Prime Western Blotting Detection Reagent (GE Healthcare Bio-Sciences, Pittsburgh, PA). Protein bands were visualized using the ChemiDoc MP Imaging System (Bio-Rad).

### RNA Extraction and Quantitative Real-time PCR

Total RNA was extracted from the tested cells using the RNeasy Mini Kit (QIAGEN, Hilden, Germany). First-strand complementary DNA (cDNA) was generated using the SuperScript™ III First-Strand Synthesis System (Invitrogen, Carlsbad, CA). Quantitative real-time PCR (qPCR) was carried out with iTaq Universal SYBR Green Master Mix (Bio-Rad, Hercules, CA) on a Bio-Rad CFX96™ Real-Time PCR Detection System. All primers were synthesized by Integrated DNA Technologies (IDT), and their sequences are provided in Table 2. The amplification protocol consisted of an initial activation step at 95 °C for 3 min, followed by 40 cycles of denaturation at 95 °C for 10 s and annealing/extension at 60 °C for 1 min. Relative mRNA expression levels were determined using the 2^−ΔΔCt method after normalization to 18s RNA or β2-microglobulin, and results were expressed as fold changes compared with the control group.

**Table 2.**
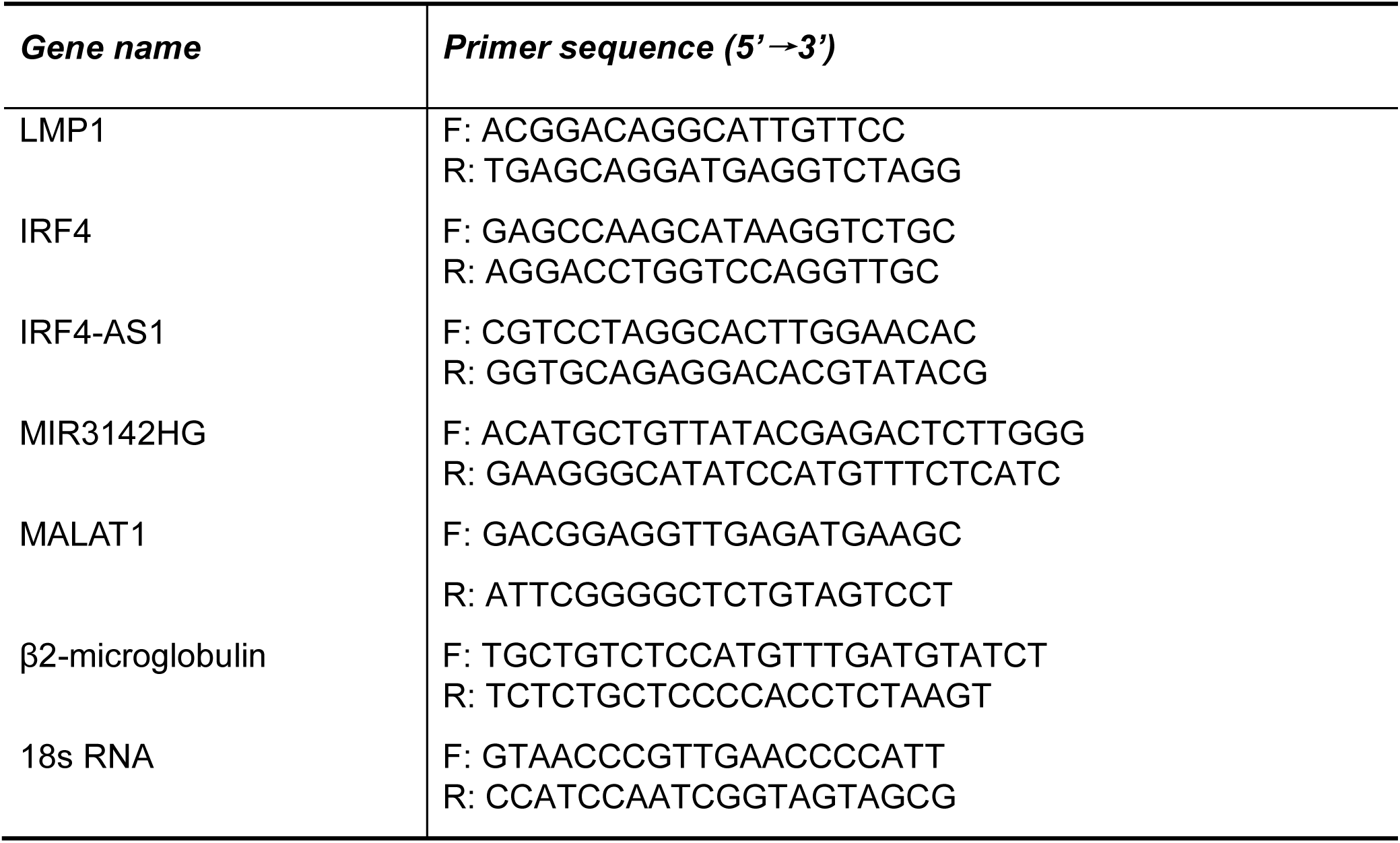
Primers for RT-qPCR.

### miRNA quantitation

For microRNA quantitation, total RNA including small RNAs was isolated using the miRNeasy Tissue/Cells Advanced Mini Kit (QIAGEN). cDNA was synthesized with the TaqMan™ Advanced miRNA cDNA Synthesis Kit (Applied Biosystems, Waltham, MA), and quantitative PCR was performed using the TaqMan™ Fast Advanced Master Mix (Applied Biosystems) together with the miR-3142 or miR-146a-5p primers/ FAM dye-labeled TaqMan MGB probe (Applied Biosystems). Expression levels of miR-3142 and miR-146a-5p were calculated using the 2^−ΔΔCt method, normalized to miR-191-5p (Applied Biosystems), and presented as fold changes relative to the control samples.

### Statistical Analysis

Statistical analyses were performed using GraphPad Prism version 10.6.0. Data are presented as the mean ± standard error of the mean (SEM). Differences among multiple experimental groups were analyzed using one-way or two-way analysis of variance (ANOVA), as appropriate for the experimental design. Comparisons between treatment and control groups were performed using paired Student’s t-tests. Correlation analyses were performed using Pearson’s or Spearman’s correlation coefficients, depending on whether the data were normally distributed. Statistical significance was defined as p < 0.05 (*), with increasing levels of significance indicated as p < 0.01 (**), p < 0.001 (***), and p < 0.0001 (****).

## Acknowledgments

This work was supported by the DoD grants HT9425-23-1-1037 (S. N.) and HT9425-25-1-0784 (L. W.), and in part by the NIH grant C06RR0306551. We are grateful to Dr. Henri-Jacques Delecluse for providing the patient-derived sLCLs and Dr. Benjamin Gewurz for providing LMP1 (and control) sgRNA plasmids. This study was conducted with the resources and facilities at the James H. Quillen Veterans Affairs Medical Center. The contents in this study are solely the responsibility of the authors and do not necessarily represent the official views of the NIH, the Department of Veterans Affairs, or the United States Government.

